# BioMedTools: a language model-powered community for biomedical computational tools

**DOI:** 10.1101/2025.05.02.651919

**Authors:** Sheng Liu, Huadong Xing, Mengying Han, Dachuan Zhang, Linlin Gong, Dongliang Liu, Junni Chen, Pengli Cai, Qian-Nan Hu

## Abstract

A large number of biomedical computational tools have spawned several tool registries. However, in the face of the rapid growth in the number of tools, existing tool registries, which are manually curated or community-driven, are difficult to keep up to date, resulting in inadequate tool repository data. In this paper, we show that language models (LMs) can aid in building a community of tools. We introduce BioMedTools (https://biomed.tools), a community of biomedical computational tools that mainly implements LM-based tool identification and a chat assistant. Compared with existing tool registries, BioMedTools achieves excellence in terms of the number of tools, frequency of data updates, and functionality. Meanwhile, the Model Context Protocol (MCP) servers hub in BioMedTools may promote the building of agents in the biomedical field. BioMedTools enables the efficient collection of tools and enhances their findability and accessibility.

## Introduction

Biomedical computational tools are computer-based tools used for research in the life sciences or medicine. The development of biomedical computational tools is advancing bioscience and medical research. According to the bioinformatics software description model biotoolsSchema, tools are described as application software with clear data processing functions. Common tools mainly include command-line tools, desktop applications, web applications, web APIs, etc.^1^ With the increasing number of biomedical computational tools, many tool registries or directories for finding and accessing scientific tools have emerged (Supplementary Table 1). Representative examples include OMICtools^2^ and bio.tools^3^. The former is a manually annotated high-quality metadata database that provides an overview of a large number of omics-related web-accessible tools. The latter is a comprehensive registry of software and databases in the life sciences based on community-driven curation. These tool databases aid the development and diffusion of scientific research tools.

However, maintaining, expanding and keeping these tool databases up to date requires significant human and financial resources. OMICtools, which attracts more than 1.5 million visitors annually, closed operations in 2020 because of a lack of financial support. The currently active biomedical computational tool library bio.tools has a staggering 30,395 tools so far. With the generation of massive biomedical big data, the creation of biomedical computational tools is also accelerating. By simply relying on the community to manually submit tool information, the latest tool progress cannot be fully captured in a timely manner. The current biomedical computational tool library suffers from imperfect data and untimely updates, which may be detrimental to researchers in finding and selecting tools. Among the 3,900 papers surveyed by Wadi et al. in 2015, 67% cited outdated software for pathway enrichment analysis^4^, which may have lead to erroneous scientific conclusions. Moreover, the explosion of tools has imposed a burden on researchers in selecting and utilizing appropriate resources. Since most researchers in the biomedical field lack a background in computer science, mastering new bioinformatics tools or databases requires a significant investment of time and effort in learning.

Another challenge posed by the explosion of tools is the increasing complexity of workflow design. As biomedical research increasingly relies on computational analyses, robust workflows or pipelines have become essential for ensuring reproducibility, scalability, and shareability of results^5, 6^. However, designing workflows generally demands specialized expertise in both biomedical and computational sciences, and the entire process requires careful curation. With the emergence of numerous complex and heterogeneous tools, workflow design has become increasingly intricate^7^. In fact, tools such as web servers are often incompatible with traditional workflows. Therefore, novel approaches to workflow construction may be necessary to address these challenges.

Recently, language models have advanced to the large language model (LLM) stage^8^. Language models aim to predict the probability of a given word sequence and are applied to text classification, entity recognition, text generation, etc. Common language models include BERT^9^, GPT3^10^ and LLaMA^11^, which are also known as foundation models. Among them, typical ones are language models based on the Transformer architecture^12^, such as the BERT and GPT series. Pre-trained language models represented by BioBERT^13^ perform well on various biomedical text mining tasks. Therefore, mining biomedical computational tool information from the literature using language models is a promising way to cope with the limitations of traditional community-based and manual curation.

In addition, language models can enhance their task-solving capabilities by invoking external tools^14, 15^. This is particularly crucial for addressing the limitations of large language models, especially in acquiring accurate information and performing scientific tasks. Therefore, biomedical artificial intelligence (AI) agents based on large language models have demonstrated significant potential in the biomedical domain by using external tools to take actions^16, 17^. Biomedical AI agents refer to specialized AI systems designed for biomedical applications, capable of engaging in scientific research and discovery much like human scientists^18^. These agents must decompose complex tasks in biomedical research into simpler subtasks and rely on a range of external tools for execution. The Model Context Protocol (MCP) provides a standardized interface for interaction between LLMs and external tools^19^, which is expected to accelerate this process. As such, LLM-based biomedical AI agents are poised to reshape bioinformatics workflows.

We aimed to develop a new LM-based method to construct a community of biomedical computational tools in the AI era. Specifically, we built BioMedTools, an AI-driven discovery platform for biomedical computational tools. It uses an LM-based tool identification system to automatically identify tools from publication titles and abstracts to achieve automatic inclusion of tools and alleviate the problem of incomplete tool data. To facilitate tool discovery and selection by biomedical researchers, we developed a web-based application that enables efficient searching and browsing of relevant tools. At the same time, BioMedTools provides rich data visualization for tool analysis and exploration. To promote a friendly user experience, we used an open-source LLM combined with a agentic Retrieval-Augmented Generation (RAG) strategy to build a question-and-answer assistant for tool information to provide a more friendly user experience. Taking a step further, in order to take advantage of the tool-use capabilities of LLM-based agents, we also developed a MCP servers collection module to promote the building of agents in the biomedical field. Overall, BioMedTools is an efficient solution for the rapid collection of tools and achieving their findability and accessibility.

## Results

### Overview of the BioMedTools community

BioMedTools aims to build a sustainable community of biomedical computational tools based on language models. It mainly includes the acquisition of tool data and the construction of an online platform. To identify potential biomedical computational tools from original research articles, we developed LM-ToolHunter, a language model-based framework for automatically identifying tools. Meanwhile, we integrated diverse data resources, including bio.tools^3^, Semantic Scholar^20^, GitHub, EDAM^21^, etc. (Supplementary Table 2) to annotate the tools from multiple perspectives.

We then leveraged these annotated datasets to construct the BioMedTools platform, which provides user-friendly access to detailed tool information. Users can not only access the tool information through regular search and browse methods, but also use the chat assistant to interact with BioMedTools. At the same time, BioMedTools provides data visualization for easy analysis of the tool.

In addition, the BioMedTools MCP servers hub allows for the collection and accumulation of MCP server for biomedical computational tools, creating a foundation of tools for intelligent agents in the biomedical domain based on large language models. Recognizing that our initial data stems from model predictions and may include inaccuracies, we have implemented a dedicated data correction system in BioMedTools. Through contributions from the community and domain expert, BioMedTools enables iteration of data and further improvement of models.

### LM-based tool identification framework

To address the limitations of existing methods for including tools in biomedical computational tool registries, we propose a language model-based tool identification framework, LM-ToolHunter (Fig. 2A).

**Fig. 1.**
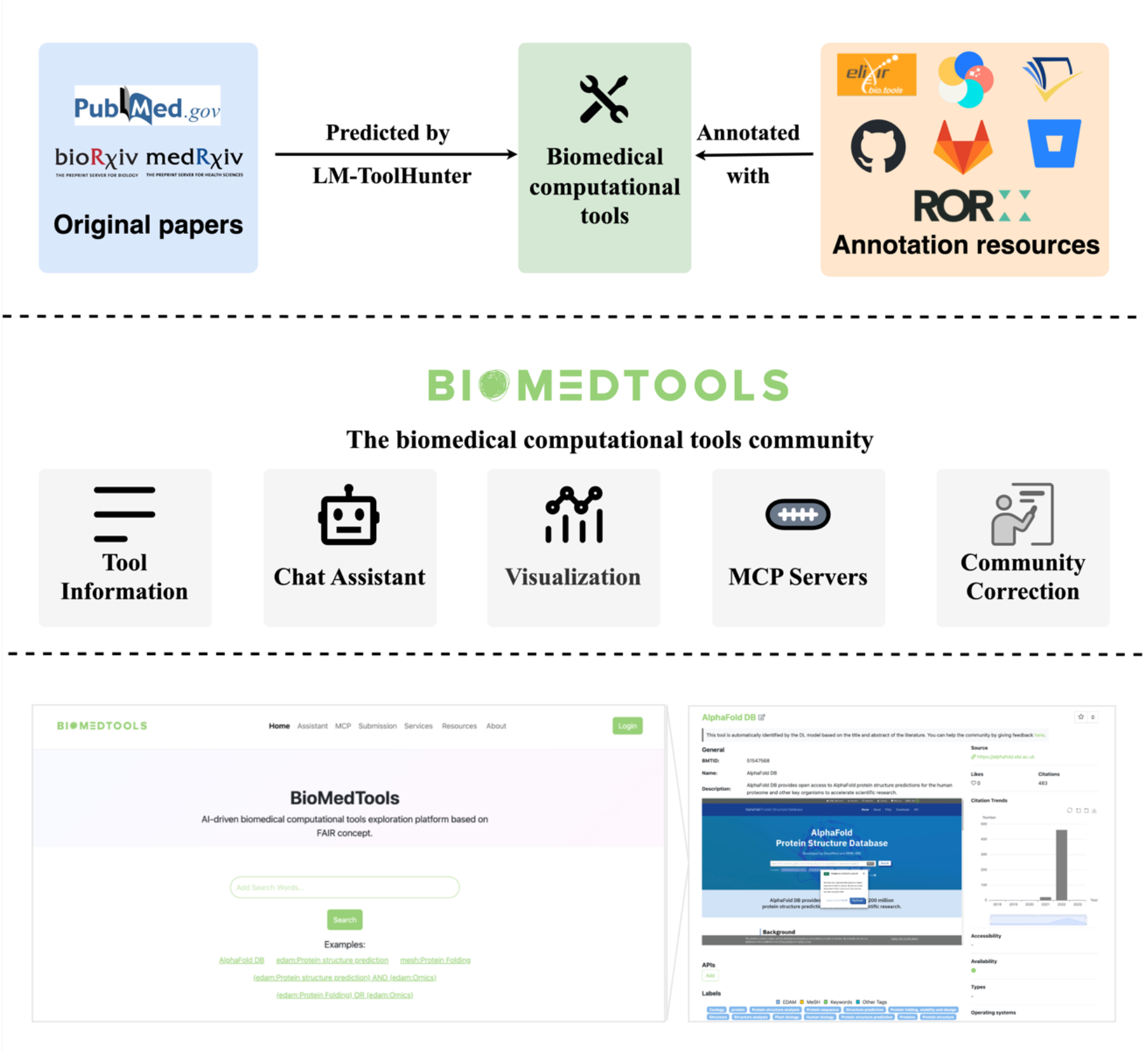
The basic workflow and main functional components of BioMedTools. First, biomedical computational tools were identified from the original papers using LM-ToolHunter from this study. Subsequently, the tools were further annotated using external public resources. The annotated tools were used to build the BioMedTools community. BioMedTools mainly includes features such as tool basic information, chat assistant, data visualization, MCP servers and community data correction.

**Fig. 2.**
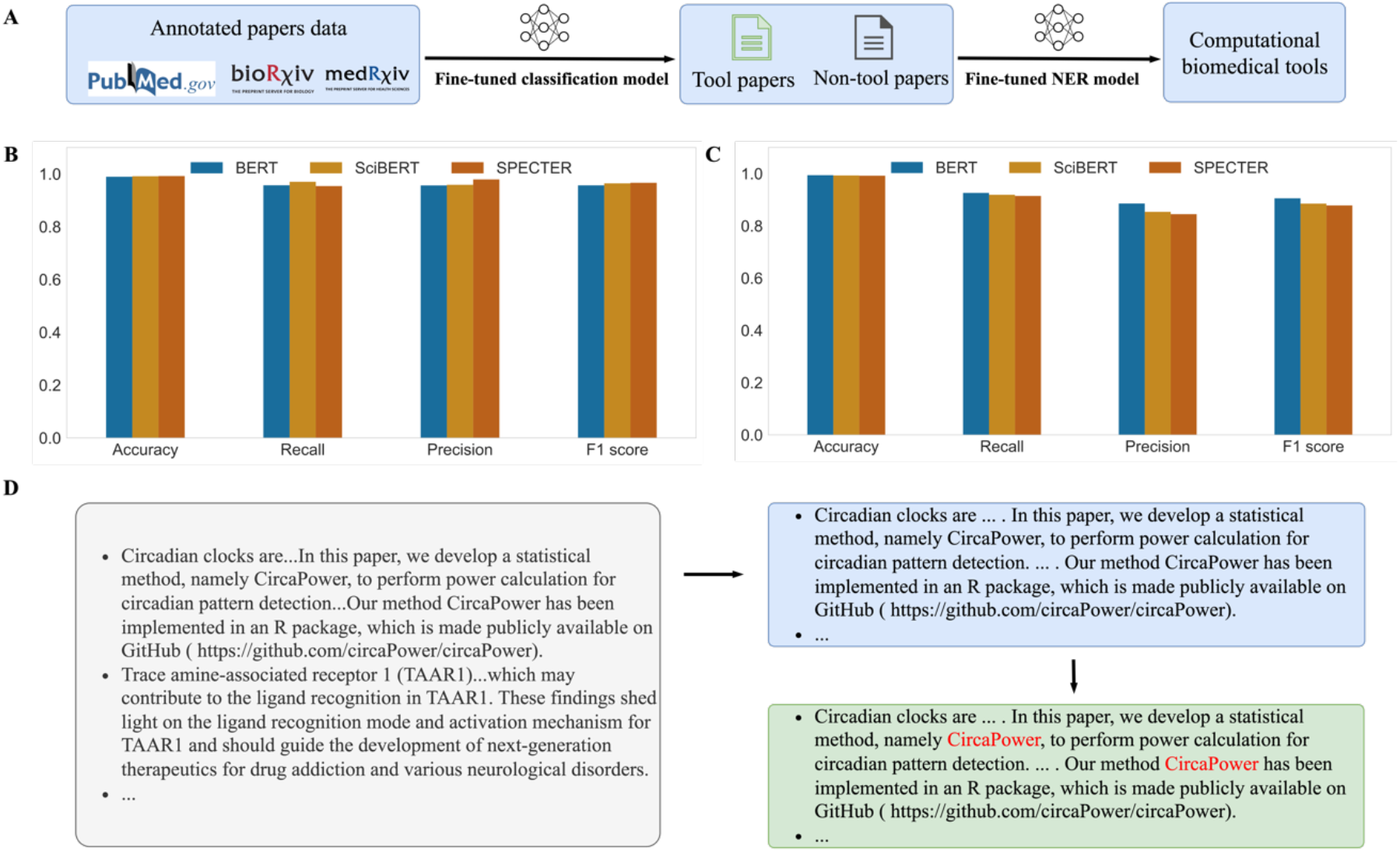
Tool identification framework LM-ToolHunter. **A** Implementation of LM-ToolHunter framework. **B** Performance of tool literature binary classification models fine-tuned on different pre-training models. **C** Performance of tool NER models fine-tuned on different pretraining models. **D** Practical working example of LM-ToolHunter.

The framework comprises two primary components. First, a tool paper classification model is used to filter articles that describe biomedical computational tools. Second, a tool entity identification model extracts tool names from the titles and abstracts of these articles. We used the dataset constructed in this study to fine-tune the pre-trained language model for different tasks and evaluate the performance of the model on a test set, respectively.

The performance metrics of the models show that for the classification model, the results of fine-tuning based on the three common pre-trained language models are excellent, with the best performance on spector, with an F1 of 0.966 (Fig. 2B). Additional evaluation metrics, including AUROC (Area Under the Receiver Operating Characteristic Curve) and AUPRC (Area Under the Precision-Recall Curve), are provided in the Appendix (Supplementary Table 3 and Supplementary Figure 1).

Meanwhile, the tool entity recognition models also performed strongly, with the best model reaching 0.905 overall (Fig. 2C and Supplementary Table 4). Ultimately, the top performing models were used for tool identification. Specifically, LM-ToolHunter first determines whether a piece of literature in the biomedical domain belongs to the biomedical computational tool literature; if it does, the system further extracts the specific tool name (Fig. 2D).

### Data visualization in BioMedTools

To analyze the trends of biomedical computational tools from multiple perspectives, we provide several visualization modules in BioMedTools. The first one is the global distribution map of tools, which allows the user to view the distribution of the number of biomedical computational tools in different scopes, Fig. 3A shows its initial interface in the web page. Each small bubble in the map corresponds to an institution, with darker colors representing a higher number of tools published by that institution. As can be seen from the map, there are significantly more research institutions in developed regions such as Europe and the United States that publish a higher number of tools than in other regions. The second one is a graph of the trend of change in the number of citations of different tools (Fig. 3B), in which the user can customize the comparison and gain insights into emerging trends in tool usage. For instance, when comparing the widely used alignment programs TopHat2^22^ and HISAT2^23^, the graph reveals that HISAT2 surpassed TopHat2 in citation count around 2019—demonstrating its rapidly growing competitiveness. In conclusion, these two visualization functions provided by BioMedTools can not only assess the situation of biomedical computational tools in different countries or institutions from a macroscopic point of view, but also compare the citation situation of different tools from a microscopic point of view.

**Fig. 3.**
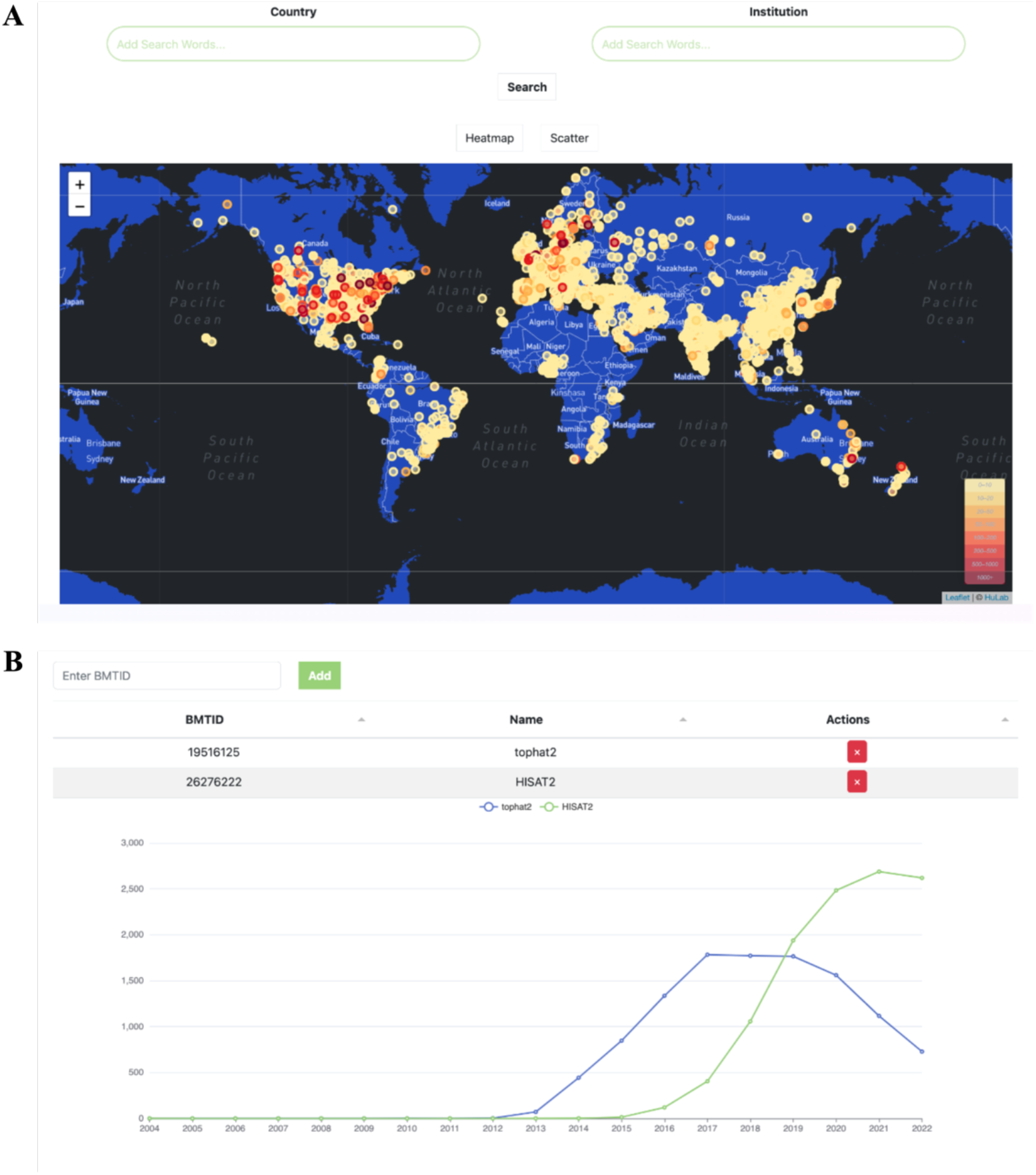
The main visualization module in BioMedTools. **A** The global map visualization for number of biomedical computational tools. **B** Curve of changes in the number of citations for alignment program TopHat2 and HISAT2.

### Chat with BioMedTools

BioMedTools features a chat-based assistant module that enables users to access information on biomedical computational tools through natural language dialogue. The core principle of this function is RAG, which operates in two main stages (Fig. 4A). The first is the construction of a vector database, in which the information text of biomedical computational tools is converted into vectors and stored in the database as a data source for retrieval. Secondly, we created the RAG-based tool conversation module, and the gray shaded part of Fig. 4A shows the specific steps. The user’s conversation history and new questions are modeled by the large language model to generate a single question, which is later used to perform a similarity search in the vector database to get relevant content blocks. The generated question and the relevant content obtained through the search are finally inputted into the large language model to realize the generation of the answer. Fig. 4B illustrates a case study in which a user engages in a natural language conversation to seek information about sequence alignment tools. This example demonstrates the chat assistant’s accuracy and its user-friendly interaction.

**Fig. 4.**
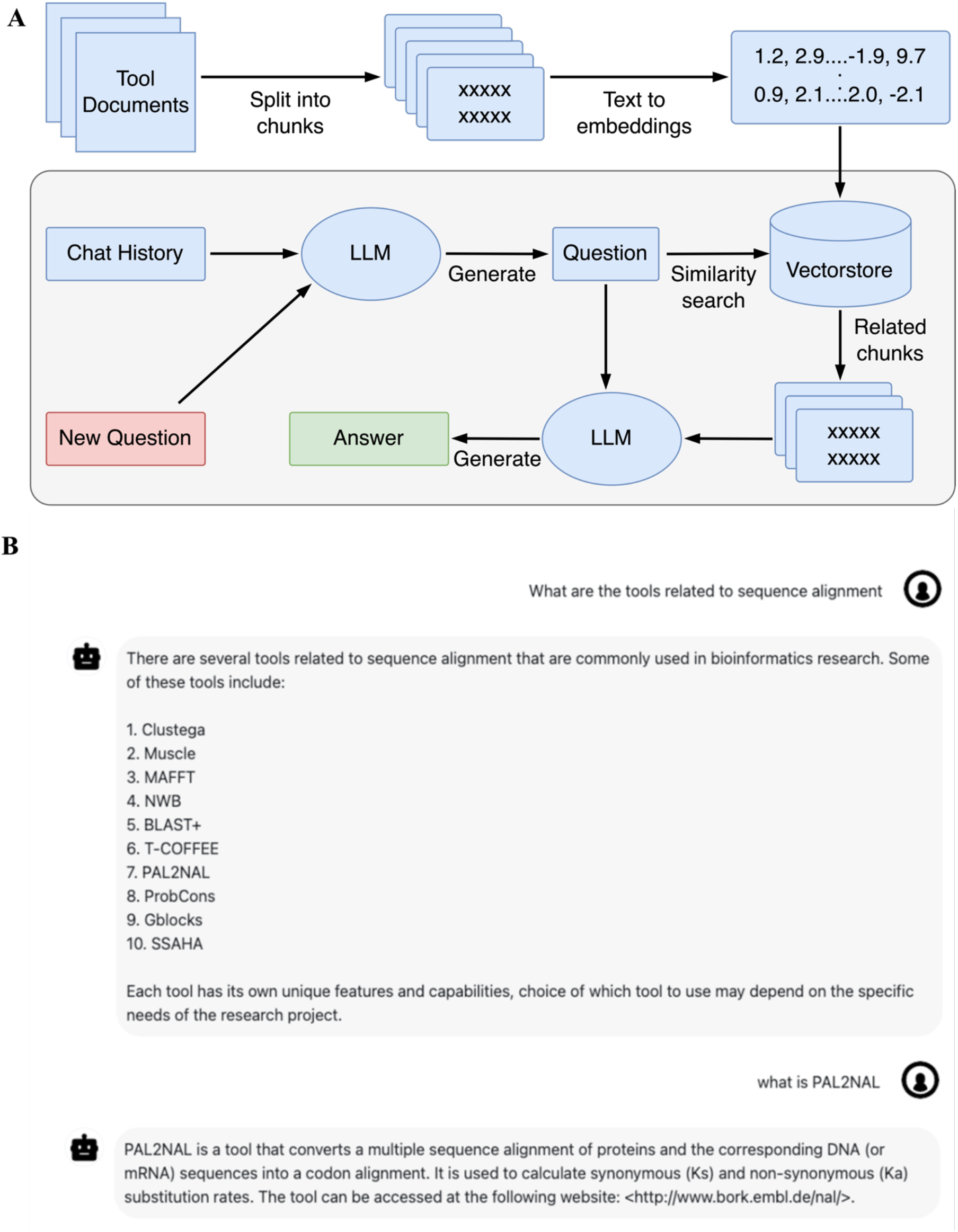
Chat assistant in BioMedTools. **A** Diagram of BioMedTools chat assistant building process. **B** A case study on the use of the BioMedTools chat assistant to consult about sequence alignment tools.

### MCP servers hub for biomedical agent building

To facilitate the development of biomedical AI agents, we have established an MCP servers hub within the BioMedTools platform, which enables the submission, download, sharing, and application of MCP servers. Users can submit biomedical MCP servers either through the tool details page or directly within the MCP section of BioMedTools.

To demonstrate the feasibility of building intelligent agents in the biomedical field based on the MCP servers provided by BioMedTools, this study initially constructed a biosynthesis agent based on a large language model using the main MCP tools of MCF2Chem^24^, AddictedChem^25^, and an unreleased pathway design tool (Fig. 5A). It can perform data queries and calculations in biosynthesis studies, such as querying the highest production titer reported in the literature for a compound or designing a biosynthetic pathway for a target compound. As shown in Fig. 5B, the biosynthesis agent correctly answered the question regarding the maximum production titer of artemisinic acid in the current MCF2Chem system. In the pathway design for artemisinic acid, the agent also reproduces the known three-step pathway for synthesizing artemisinic acid from acacia diene. In contrast, when the same query was posed to GPT-4, it was able to identify the biosynthetic pathway but failed to correctly provide the compound Simplified Molecular Input Line Entry System (SMILES), highlighting the necessity of external tools for rigorous and precise factual content. Interestingly, the intelligent agent was not provided with the corresponding compound SMILES in the dialog, and it automatically selected and executed the tool for obtaining the SMILES for pathway design.

**Fig. 5.**
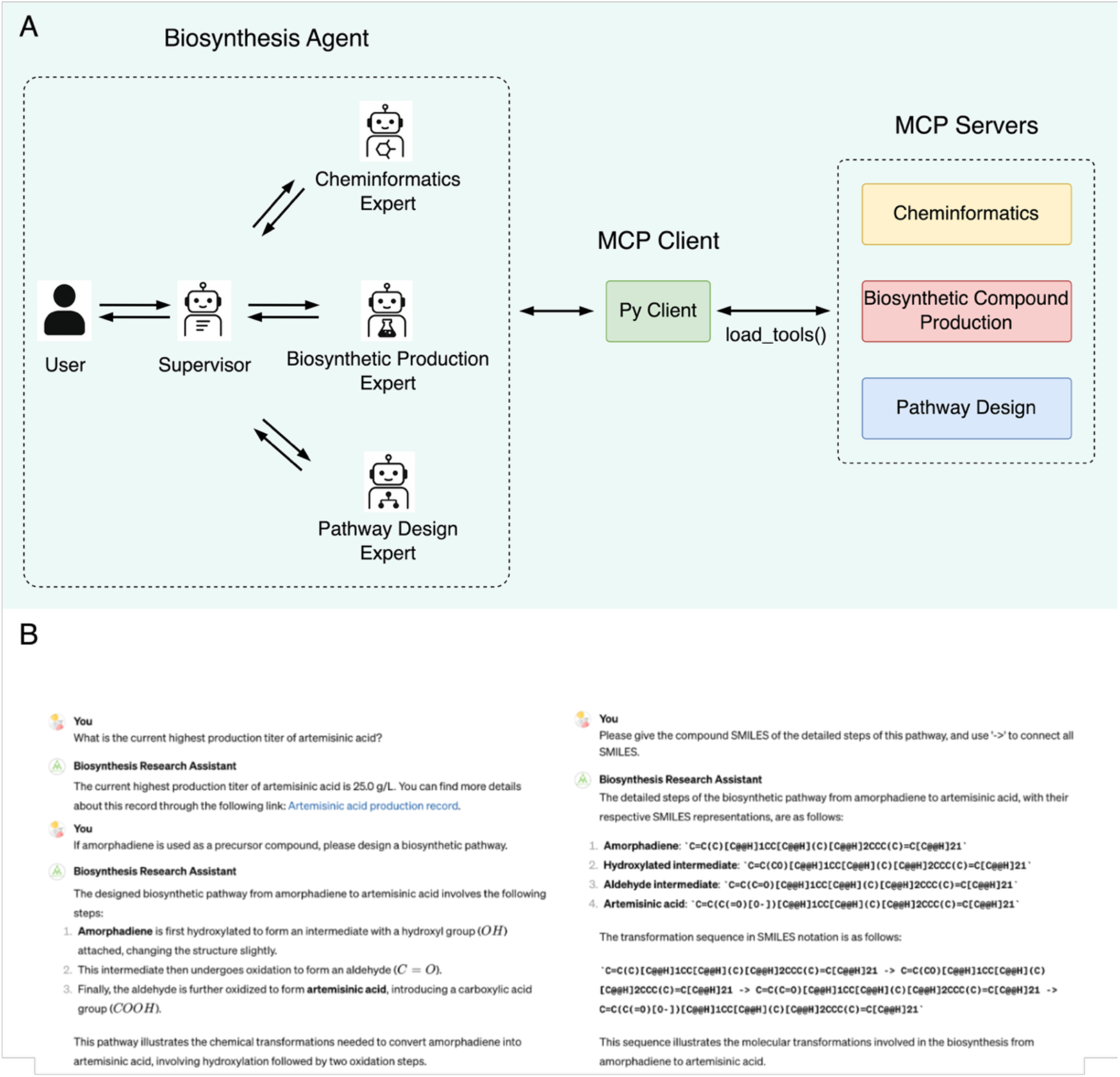
Biosynthesis agent building and application case demonstration. **A** The overall framework of the biosynthesis agent based on LLM and MCP servers. The agent is built on Qwen3-32B and implemented using the LangGraph framework. **B** Use biosynthesis agent to query artemisinic acid production information and design pathway. In order to make the content more compact, this picture removes blanks and other irrelevant parts based on the original screenshot.

### Comparison with similar platform

The main purpose of the biomedical computational tool registry is to facilitate the findability and accessibility of tools. To illustrate the strengths of BioMedTools in this regard, we conducted a comprehensive comparison with the major tool repositories in existence (Tab.1). For tool findability, BioMedTools includes 493,647 tools—far more than bio.tools and Database Commons—and, like bio.tools, it covers almost all tool types. Unlike current tool registries that rely on manual curation or community contributions for data updates, BioMedTools integrates AI and community engagement to enable continuous and efficient data updating. Without human intervention, BioMedTools can achieve daily collection of basic information on tools, meeting the needs of researchers to access the latest tools. To realize higher quality data, BioMedTools also provides a community-driven data update system. In terms of user interaction, both bio.tools and BioMedTools provide APIs in addition to regular web-based searching and browsing. With the rapid advancement of LLMs, BioMedTools is designed to support future development. Compared to existing tool platforms, it provides an AI chat assistant for accessing tool information. In addition, the MCP servers hub in BioMedTools offers a convenient entry point for the biomedical computational tool community to discover available MCP servers.

**Table 1.** Comparison of BioMedTools with other common tool platforms (As of April 29, 2025)

| Platform | Number of tools | Tool types | Availability status | Data update method | Frequency of data update | AI chat assistant | API | MCP servers |
| --- | --- | --- | --- | --- | --- | --- | --- | --- |
| bio.tools | 30,395 | **all types** | no | community driven | irregularly | no | **yes** | no |
| Database Commons | 6,961 | only database | **yes** | manual curated/community driven | irregularly | no | no | no |
| BioMedTools | **493,647** | **all types** | **yes** | **AI automated/community driven** | **everyday** | **yes** | **yes** | **yes** |

## Discussion

The FAIR guidelines, which aim to improve the findability, accessibility, interoperability, and reusability of digital resources, have been widely recognized and adopted by industries such as research data and software^26^. However, current registries of biomedical computational tools suffer from imperfect data and untimely updates, and the maintenance of tool information often relies on manual hand-holding. Therefore, in this study, we developed a new approach based on language models for the construction of biomedical computational tool communities in the AI era. Specifically, we constructed BioMedTools, an AI-driven exploration platform for biomedical computational tools. It automatically discovers tools from literature titles and abstracts using a language model-based tool recognition system to automate tool inclusion and mitigate the problem of imperfect tool data. Overall, BioMedTools is an effective solution for rapid collection of tools and their findability and accessibility.

In order to realize the identification of biomedical computational tools from the literature, this study proposes a language model-based tool identification framework, LM-ToolHunter. Despite the high performance of the associated deep learning models on the test set, there are still mistakes in the actual predictions. Therefore, in order to achieve a balance between quantity and quality of tool data, this study developed a community-driven data correction system in the BioMedTools platform. In contrast to similar purely community-driven bioinformatics tool registries in bio.tools, BioMedTools provides machine prediction information for almost all tools. Users can revise and extend the information from the initial version of the tool. Theoretically, the more a tool is noticed by researchers, the more likely it is to be further annotated by humans, and BioMedTools’ concept of iterative data building based on both AI and community-driven data can effectively improve the efficiency of building biomedical computational tool data.

In addition to the acquisition of raw data, this study utilizes open-source large language models combined with RAG strategies to construct a chat assistant for tool information to provide a more user-friendly experience. With the popularity of large language model application development, this form of conversational interaction may be a future trend for bioinformatics tools. Currently, compared to fine-tuning the large language model, the RAG strategy is a better choice in the field of research tools that require higher information accuracy. Restricted by the fact that tool data are derived from machine predictions, the chat assistant in BioMedTools is not always satisfactory. However, as the data in BioMedTools continues to be corrected, this situation will improve accordingly. Also, as data from user interactions with BioMedTools accumulates, we will customize the large language model in the field of biomedical computational tools.

The current BioMedTools automatically collects literature from PubMed, bioRxiv, and medRxiv, and although it already includes most of the literature in the biomedical field, it also may omit some computational tools in fields such as biology or chemistry. Therefore, in future studies, a wider range of literature, such as arXiv and ChemRxiv, could be encompassed in order to be able to keep track of the advances in biomedical computational tools. In addition, the methodology in this study only addresses the titles and abstracts of the literature, which can overlook more detailed information about the tools. In the future, new models can be developed for extracting tool information from the full text of the literature for literature that can be accessed in full text. Comprehensive access to tool information will facilitate the comparison and analysis of tools, thus promoting the application and development of biomedical computational tools.

In order to assess the citation of the tool, this study obtained the citation count of the tools’ literature from Semantic Scholar. However, not all citations have the same effect. There have been studies that use deep learning to classify citations in order to determine whether the statement provides support or comparative evidence for the cited work, or just mentions it^27^. In the future, it will be possible to access the citation context of the tool literature and classify citations based on different citation intentions. For example, some citations are motivated by comparisons of similar tools, whereas others may have used the tool in a study. For those citations that actually used the tool, the citation context can also be used as the scholarly usage scenarios data of the tool, which can provide a data basis for providing more accurate tool recommendations, and, at the same time, a more objective assessment of the impact of a tool.

The ability of agents to use external tools can compensate for the shortcomings of large language models, especially for the acquisition of accurate scientific knowledge and the execution of scientific tasks, and there have been several studies applying agents to biomedical research^28, 29, 30^. For example, GeneGPT enables a large language model to learn to use web APIs from the National Center for Biotechnology Information (NCBI) through context learning to achieve optimal performance on the GeneTuring dataset^30^. Considering the potential application of large language model-based agents in specialized research areas, this study constructed a MCP servers hub. Community users can submit, download, and cite MCP servers through the BioMedTools online platform. In the future, these MCP servers will be available as a resource for agent research and applications in the biomedical field. Biomedical AI Agent based on the combination of large language models and MCP servers may become a new paradigm for building biomedical computational workflows.

## Methods

### Data sources, managing and updating

This study identifies and includes biomedical computational tools from literature big data along with a comprehensive informational annotation of the tools, thus involving several different data sources. Among them, literature is the most dominant data, and they include the biomedical literature database PubMed, the preprint server bioRxiv, and medRxiv. Literature from the above three data sources is used for the identification of biomedical computational tools as well as for the annotation of the tools.

In addition to this, all other data sources were used for the comprehensive annotation of the tools. For example, the citation data of the tools is obtained from Semantic Scholar, as it has more than 200 million documents and supports bulk download method. The registries bio.tools^3^ and Database Commons^31^ for tools in the biomedical field, as well as the code repositories GitHub, GitLab, and Bitbucket were used for multidimensional annotation of the tools. Meanwhile, this study uses data from Research organizations registry (ROR) to normalize the affiliation of the tools. The EDAM ontology data is then used for the prediction of EDAM terms. For links to the data sources in this study, refer to the list of data sources in the supplementary materials (Supplementary Table 2). All data were parsed into JSON format and stored in MongoDB after acquisition, with literature data updated regularly on a daily basis and other data updated semi-annually or annually.

### Datasets of models

In order to create a binary classification dataset of tool literature, the literature corresponding to tools in bio.tools was used as positive example data in this study, while a portion of the data was manually labeled as positive examples. Negative example data, on the other hand, were obtained from medical literature data that had a high probability of not being tool literature, as well as a portion of manually labeled literature. Each entry consists of text consisting of article titles and abstracts as well as labels of whether or not they belong to the tool literature, resulting in 16,784 positive examples and 147,098 negative examples, respectively. For the tool entity identification task, we mapped the tool names of tool-based literature back to the original text consisting of the title and abstract of the literature to get the starting and ending positions of the tool entities.

### Fine-tuning model

The classical language representation model, BERT^9^, is widely used in various natural language processing tasks. Various versions of BERT have been subsequently derived, including SciBERT^32^, a pre-trained language model for scientific texts, and BioBERT^13^, a biomedical language representation model. In addition, the SPECTER^33^ language model allows for representation learning at the scientific document level. In order to identify biomedical computational tools from biomedical literature big data, the pre-trained models described above were fine-tuned to build a text binary classification model and a tool entity recognition model. For the training of the classification model, we used the AdamW optimizer with a learning rate of 5×10-5. For the NER model, we used the Adam optimizer in combination with the OneCycleLR^34^ learning rate scheduler to dynamically adjust the learning rate and improve the training efficiency and performance of the model, with the maximum number of training iterations of the model being 100. The model was trained and evaluated using Python 3.7.10 with Pytorch 1.10.0.

### Annotation of biomedical computational tools

To provide a comprehensive annotation of biomedical computational tools, this study uses three ways to maintain the 16 tool information categories. The first approach involves automatic prediction or extraction of information through models or programs, which involves fields such as name, description, URL, and so on. The second way is to automatically integrate external data sources such as bio.tools and GitHub, which involves fields such as languages, cost, accessibility, and so on. Another way is to continuously improve the tool data through community contributions. Except for citation count and publications, all other fields can be contributed by the community and manually corrected under constraints, and these fields are known as fields that can be manually intervened. By combining these strategies and assigning appropriate priorities to each data source, BioMedTools achieves a continuously evolving, high-quality repository of tool information.

### Implementation of BioMedTools online platform

The BioMedTools was deployed with FastAPI 0.73.0 for building APIs and Vue.js 3.3.2 for building web user interfaces. We used Echarts 5.4.2 and Highcharts 11.0.1 for interactive visualization and render map with leaflet 1.2.0. All relevant data is stored in a MongoDB 5.0.4 database. To better search for tools, we have developed a powerful search function using Elasticsearch 7.16.2. For building chat assistant in BioMedTools, we first spliced the main information of all the tools in BioMedTools into text chunks and used the BAAI/bge-m3^35^ model to generate text embeddings, which were then stored in the vector database Milvus^36^. To achieve improved retrieval performance in RAG, the reranking model BAAI/bge-reranker-v2-m3^35^ is employed to optimize the initial retrieval results. The LLM used by Chat assistant is based on the open source LLM Qwen3-32B^37^ and is deployed locally using the vLLM^38^. We built the entire backend of the conversation assistant using the LangChain and LangGraph framework.

## Acknowledgements

This work was financially supported by the National Key Research and Development Program of China [grant numbers: 2020YFA0908302, 2019YFA0904300 and 2021YFC2103001]. We also extend our gratitude to Dr. Ziniu Dai from the School of Medicine, Westlake University, for his efforts in organizing several commonly used computational and analytical tools in biomedical research.

## Author contributions

S.L., H.X. and M.H. designed and conducted this study; The BioMedTools platform was validated and improved by general user feedback from D.Z., L.G., D.L., J.C. and P.C.; P.C. and Q.H. supervised the study. The original draft was written by S.L., H.X., and M.H., and all authors contributed to reviewing and editing of the final manuscript.

## Materials & Correspondence

Correspondence and requests for materials should be addressed to Pengli Cai and Qian-Nan Hu.

## Supplementary Information

**Supplementary Table 1.** Biomedical computational tool registries.

| Name | URL | Status |
| --- | --- | --- |
| DAS Service Registry | www.dasregistry.org | Unavailable |
| EMBRACE Registry | www.embraceregistry.net | Unavailable |
| BioMOBY Central | www.biomoby.org | Available |
| seekda | www.seekda.com | Unavailable |
| BioCatalogue | http://www.biocatalogue.org/ | Unavailable |
| SEQanswers wiki | http://wiki.SEQanswers.com/ | Unavailable |
| SIB bioinformatics resource portal | https://www.expasy.org/ | Available |
| OMICtools | http://omictools.com/ | Unavailable |
| NAR Database Issues | http://www.oxfordjournals.org/nar/database/c/ | Available |
| Bioinformatics Links Directory | http://bioinformatics.ca/links_directory/ | Unavailable |
| Bioconductor | http://www.bioconductor.org/ | Available |
| Bio-Linux | - | - |
| Debian Med | http://debian-med.alioth.debian.org | Unavailable |
| BioJS | http://biojs.net/ | Available |
| BioStar Tools | https://www.biostars.org/t/Tools/ | Available |
| bio.tools | https://bio.tools | Available |
| Dockstore | https://dockstore.org/ | Available |
| Aviator | https://www.ccb.uni-saarland.de/aviator | Available |
| Database Commons | http://ngdc.cncb.ac.cn/databasecommons/ | Available |

**Supplementary Table 2.** Data sources used for BioMedTools construction.

| Data source | Category | URL |
| --- | --- | --- |
| PubMed | Publication | https://pubmed.ncbi.nlm.nih.gov/download/ |
| bioRxiv | Publication | https://api.biorxiv.org/ |
| medRxiv | Publication | https://api.medrxiv.org/ |
| Semantic Scholar | Publication | https://api.semanticscholar.org/datasets/v1/release/latest |
| bio.tools | Tools registry | https://biotools.readthedocs.io/en/latest/api_reference.html |
| Database Commons | Tools registry | https://ngdc.cncb.ac.cn/databasecommons/ |
| GitHub | Tools registry | https://docs.github.com/en/rest |
| GitLab | Tools registry | https://docs.gitlab.com/ee/api/ |
| Bitbucket | Tools registry | https://developer.atlassian.com/bitbucket/api/2/reference/ |
| ROR | Research organizations registry | https://ror.org/ |
| EDAM | Ontology | https://edamontology.org/ |

**Supplementary Table 3.** Performance of different biomedical computational tool publication classification models.

| Method | TP | FP | TN | FN | Precision (%) | BA (%) | F1 score (%) | MCC (%) |
| --- | --- | --- | --- | --- | --- | --- | --- | --- |
| BERT::title+abstract | 1,610 | 73 | 12,046 | 72 | 95.66% | 97.56% | 95.69% | 95.09% |
| SciBERT::title+abstract | 1,632 | 70 | 12,049 | 50 | 95.89% | 98.22% | 96.45% | 95.96% |
| SPECTER::title+abstract | 1,605 | 34 | 12,085 | 77 | 97.93% | 97.57% | 96.66% | 96.21% |

**Supplementary Table 4.** Overall performance of the named entity recognition (NER) model for biomedical computational tools.

| Method | Precision (%) | Recall (%) | F1 score (%) | Accuracy (%) |
| --- | --- | --- | --- | --- |
| BERT::title+abstract | 88.58% | 92.61% | 90.55% | 99.40% |
| SciBERT::title+abstract | 85.39% | 91.92% | 88.53% | 99.29% |
| SPECTER::title+abstract | 84.50% | 91.45% | 87.84% | 99.20% |

**Supplementary Figure 1.**
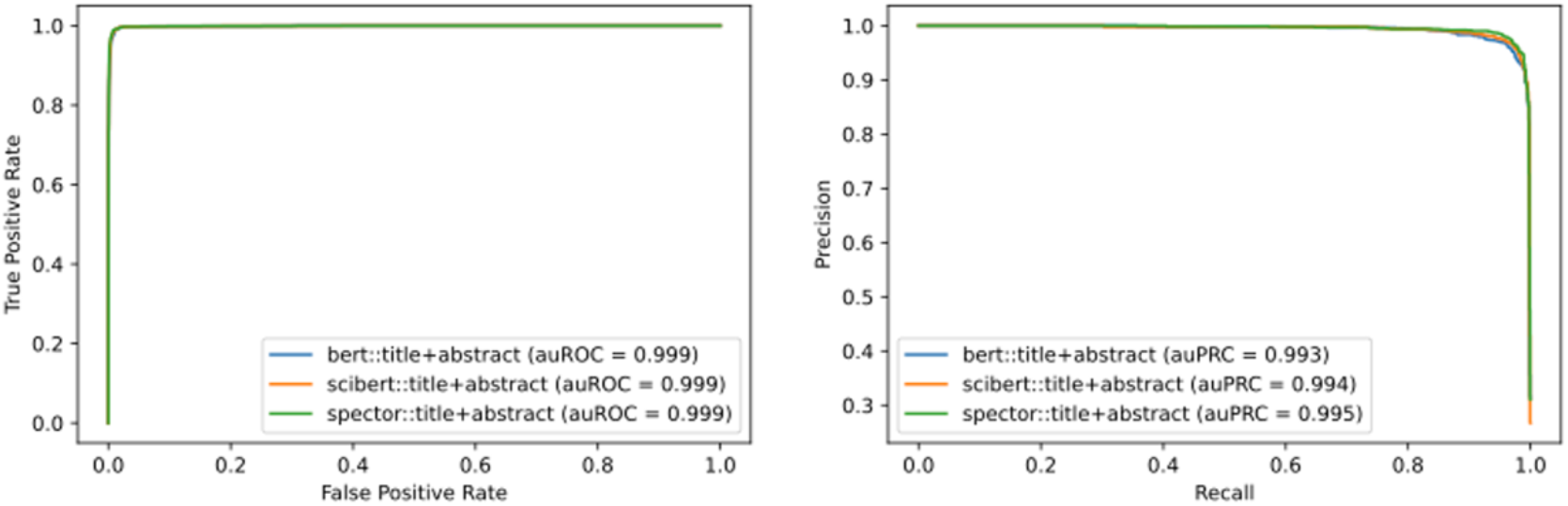
Area under the receiver operating characteristic curves (AUROCs) for the biomedical computational tool publication classification models (left). Area under the precision-recall curves (AUPRCs) for the biomedical computational tool publication classification models (right).

